# Trends of *Plasmodium falciparum* molecular markers associated with resistance to artemisinins and reduced susceptibility to lumefantrine in Mainland Tanzania from 2016 to 2021

**DOI:** 10.1101/2024.01.18.576107

**Authors:** Catherine Bakari, Celine I. Mandara, Rashid A. Madebe, Misago D. Seth, Billy Ngasala, Erasmus Kamugisha, Maimuna Ahmed, Filbert Francis, Samwel Bushukatale, Mercy Chiduo, Twilumba Makene, Abdunoor M. Kabanywanyi, Muhidin K. Mahende, Reginald A. Kavishe, Florida Muro, Sigsbert Mkude, Renata Mandike, Fabrizio Molteni, Frank Chacky, Dunstan R. Bishanga, Ritha J. A. Njau, Marian Warsame, Bilali Kabula, Ssanyu S. Nyinondi, Naomi W. Lucchi, Eldin Talundzic, Meera Venkatesan, Leah F. Moriarty, Naomi Serbantez, Chonge Kitojo, Erik J. Reaves, Eric S. Halsey, Ally Mohamed, Venkatachalam Udhayakumar, Deus S. Ishengoma

## Abstract

**Background:** Therapeutic efficacy studies (TESs) and detection of molecular markers of drug resistance are recommended by the World Health Organization (WHO) to monitor the efficacy of artemisinin combination therapy (ACT). This study assessed the trends of molecular markers of artemisinin resistance and/or reduced susceptibility to lumefantrine using samples collected in TES conducted in Mainland Tanzania from 2016 to 2021.

**Methods:** A total of 2,015 samples were collected during TES of artemether-lumefantrine at eight sentinel sites (in Kigoma, Mbeya, Morogoro, Mtwara, Mwanza, Pwani, Tabora, and Tanga regions) between 2016 and 2021. Photo-induced electron transfer polymerase chain reaction (PET-PCR) was used to confirm presence of malaria parasites before capillary sequencing, which targeted two genes: *Plasmodium falciparum* kelch 13 propeller domain (*k13*) and *P. falciparum* multidrug resistance 1 (*pfmdr1*).

**Results:** Sequencing success was ≥87.8%, and 1,724/1,769 (97.5%) *k13* wild-type samples were detected. Thirty-seven (2.1%) samples had synonymous mutations and only eight (0.4%) had non-synonymous mutations in the *k13* gene; seven of these were not validated by WHO as molecular markers of resistance (I416V, E433D, R471S, P475S, A578S, and Q613E). One sample from Morogoro in 2020 had a *k13* R622**I** mutation, which is a validated marker of artemisinin partial resistance. For *pfmdr1,* all except two samples carried N86 (wild-type), while mutations at Y184**F** increased from 33.9% in 2016 to about 60.5% in 2021, and only four samples (0.2%) had D1246**Y** mutations. *pfmdr1* haplotypes were reported in 1,711 samples, with 985 (57.6%) NYD, 720 (42.1%) N**F**D, and six (0.4%) carrying minor haplotypes (three with NY**Y**, 0.2%; Y**F**D in two, 0.1%; and N**FY** in one sample, 0.1%). Between 2016 and 2021, NYD decreased from 66.1% to 45.2%, while N**F**D increased from 38.5% to 54.7%.

**Conclusion:** This is the first report of the R622I (k13 validated mutation) in Tanzania. N86 and D1246 were nearly fixed, while increases in Y184**F** mutations and N**F**D haplotype were observed between 2016 and 2021. Despite the reports of ART-R in Rwanda and Uganda, this study did not report any other validated mutations in these study sites in Tanzania apart from R622I suggesting that intensified surveillance is urgently needed to monitor trends of drug resistance markers and their impact on the performance of ACTs.

## Background

Antimalarial drugs, particularly artemisinin-based combination therapy (ACT), are recommended and widely used for effective case management, but drug resistance is a major threat that has impacted their effectiveness for malaria control and elimination. The threat is higher, especially in sub-Saharan African (SSA) countries, which contributed over 95.0% of cases and deaths globally in 2021 [1]. ACTs were introduced in the early 2000s in most malaria-endemic countries following the World Health Organization’s (WHO) recommendations due to widespread resistance to previously used antimalarials, including chloroquine (CQ) and sulphadoxine-pyrimethamine (SP) [2]. Before and after adoption of ACTs, their efficacy has remained above 90.0% in many countries in SSA, including Tanzania, and this has played a vital role in the reduction of malaria burden between 2000 and 2015 [1]. However, progress has stalled since 2015, and the emergence of artemisinin partial resistance (ART-R) in Africa threatens the gains attained over the past two decades and the ongoing elimination efforts [1]. Thus, there is an urgent need to intensify surveillance to monitor the efficacy as well as track the emergence and spread of antimalarial-resistant parasites, particularly in SSA [3].

Following deployment of ACTs, WHO recommended monitoring both the efficacy and safety of ACTs to support effective case management strategies [4]. According to WHO, analysis of molecular markers associated with antimalarial resistance should also be done within TES to capture the emergence and track the spread of resistant parasites to artemisinins and partner drugs. Polymorphisms in different parasite genes, including *Plasmodium falciparum* kelch 13 (*k13*), *Plasmodium falciparum* chloroquine resistance transpo*rter (pfcrt), Plasmodium falciparum* dihydrofolate reductase *(pfdhfr), Plasmodium falciparum* dihydropteroate synthase *(pfdhps),* and *Plasmodium falciparum* multidrug resistance 1 *(pfmdr1),* have been identified and are commonly used as key molecular markers for tracking antimalarial resistance to the respective drugs [5,6]. For commonly used ACTs such as artemether-lumefantrine (AL), mutations in *k13* have been linked to ART-R while mutations in *pfmdr1* are associated with tolerance or reduced sensitivity to lumefantrine and/or resistance to other drugs such as CQ and amodiaquine (AQ). Currently, WHO recommends monitoring any of the 13 validated non-synonymous mutations in *k13* as markers of ART-R(A675**V**, R622**I**, C580**Y**, P574**L**, R561**H,** P553**L,** I543**T**, R539**T**, Y493**H**, M476**I**, C469**Y**, N458**Y**, and M476**I**), while eight mutations are considered to be candidate markers (P441**L**, G449**A**, C469**F**, A481**V**, R515**K**, P527**H**, G538**V**, and V568**G**) [7]. In recent years, ART-R has been confirmed in multiple African countries, particularly in the Horn of Africa in Eritrea [8] associated with the R622**I.** Reports have also come from East African countries like: 1) Rwanda, where ART-R was associated with R561**H** mutation [9]; 2) Tanzania, with R561**H** and 675 mutations [10,11] and 3) Uganda, where C469**Y** and A675**V** mutations have been reported [11, 12]. Due to the threat of potential spread and impact of ART-R in Africa, monitoring the efficacy of current and future antimalarials through clinical evaluation and detection of drug resistance markers is necessary and urgently needed.

The commonly detected polymorphisms within the *pfmdr1* gene include N86**Y**, Y184**F**, S1034**C**, N1042**D,** and D1246**Y,** and the most frequently reported mutations are N86**Y**, Y184**F**, and D1246**Y** [5]. Whereas N86 (wild-type) has been linked to reduced sensitivity to lumefantrine, the 86**Y** mutation has been associated with reduced sensitivity and/or resistance to AQ and CQ. The presence of the N86, 184**F**, and D1246 (N**F**D) haplotypes is linked to reduced sensitivity to lumefantrine, while the 86**Y**, Y184, and 1246**Y** (**Y**Y**Y**) haplotypes have been reported to cause decreased sensitivity to CQ and AQ [12]. However, the mutations in the *pfmdr1* gene have not been associated with clinical failure or resistance to lumefantrine, and there is no recommended marker for this important partner drug.

In Tanzania, TES for ACTs has been implemented before and after they were deployed as recommended by WHO to ensure effective case management [13–16]. After the deployment of ACTs in 2006 [18, 19], TES have normally focused on AL, which is the first-line antimalarial drug for the treatment of uncomplicated falciparum malaria [15, 20], together with alternative ACTs such as artesunate-amodiaquine (ASAQ), the first-line drug in Zanzibar since 2003 [21, 22], and dihydroartemisinin-piperaquine (DP), which was deployed in Mainland Tanzania in 2014 [17]. These TESs have been implemented by the Technical Working Group (TWG) of the National Malaria Control Programme (NMCP) and its partners, and the studies have been consistently done since 1997. Together with the studies done in the 2000s, the TES have shown that AL has maintained a cure rate of over 95.0%, while the cure rate of DP has been reported to be >98.0% [14,18]. The efficacy of ASAQ was less than 90.0% before the deployment of ACTs in 2006 [15], but its performance has increased, reaching 100% in 2017, possibly linked to the withdrawal of CQ in Tanzania [16, 17].

Like in many malaria-endemic countries, routine detection of molecular markers of antimalarial resistance was not consistently performed as part of TES in Tanzania due to limited capacity. In 2016, the capacity to detect markers of resistance to different drugs was established with the support of the Partnership for Antimalarial Resistance Monitoring in Africa (PARMA) network [24], and the analysis has been performed as part of TES except in 2017 which is not covered in this study. The molecular analysis that has been done within TES since 2016 aimed at generating data on mutations in *pfmdr1* and *k13.* In 2021 ART-R was detected in Tanzania, through country-wide surveys of malaria parasites [10] and confirmed using TES [11]. Thus, it is critical to conduct a thorough assessment of markers of drug resistance in Tanzania using retrospective and prospectively collected samples and data to explore the presence of ART-R in the sites covered by TES. This study was undertaken using the data generated through molecular analysis of samples collected in TESs from 2016 to 2021 to assess the trends of markers of resistance to artemisinins (*k13*) and/or reduced susceptibility to lumefantrine (*pfmdr1*). The findings provide evidence to NMCP and its partners to support malaria case management guidelines and policy as well as identification of foci of ART-R and planning of targeted malaria molecular surveillance (MMS) with a focus on areas with high prevalence and risk of resistant parasites in Mainland Tanzania.

## Methods

### Study site

This study was based on the analysis of samples that were collected by the TWG of NMCP during TESs, which were conducted to assess the efficacy and safety of AL. The samples were collected from 2016 to 2021 at eight TES sentinel sites of NMCP, which were: Igombe health centre (Mwanza), Ipinda (Mbeya), Yombo (Pwani), Mkuzi (Tanga), Mlimba (Morogoro), Nagaga (Mtwara), Simbo (Tabora) and Ujiji health centre (Kigoma) (Figure 1). The sites and details of study design and sample collection have been fully described in previous TES [18] and were based on the WHO protocol of 2009 [4]. According to the TWG’s framework, each sentinel site conducts a TES at least once every two years and thus all the sites were sampled three times during the study period. However, TES 2017 samples were not genotyped and are not included in this study.

**Figure 1:**
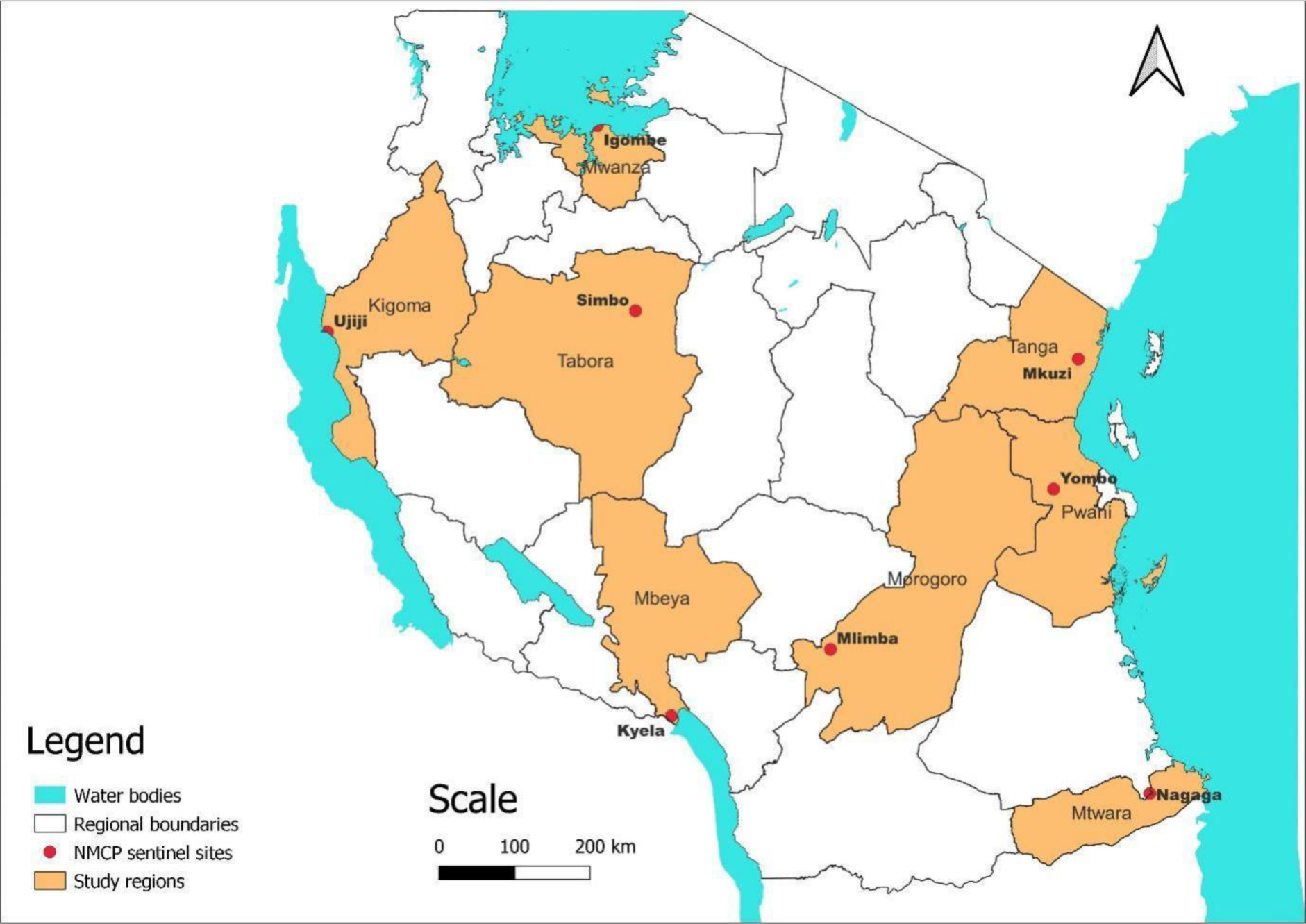
Map of Tanzania showing the regions and the eight National Malaria Control Programme sentinel sites.

### Sample collection, processing and molecular analysis

The studies that provided samples for this analysis enrolled malaria patients meeting specific criteria as per the WHO protocol of 2009 [4], and details of enrolment procedures have been fully described in a previous study [18]. In summary, enrolled patients were aged 6 months to 10 years and had uncomplicated malaria with *P. falciparum* mono-infections and 250–200,000 asexual parasites/µl of blood, as well as fever at presentation or history of fever in the past 24 hours. The enrolled patients received AL and were followed up for 28 days as per WHO protocol [4].

Dried blood spot (DBS) samples were collected on Whatman 3-mm filter paper (Whatmann No. 3, GE Healthcare Life Sciences, PA, USA) from enrolled patients on Day 0 (before treatment) and during follow-up visits in the case of recurrent infections, according to the procedures described earlier [18]. Genomic DNA was extracted using QIAamp blood mini-kits (Qiagen GmbH, Hilden, Germany) according to the manufacturer’s instructions and stored at 4°C before use. Molecular analysis was performed on all samples collected upon enrolment (day 0) and during follow-up in the case of recurrent infections to confirm the presence of malaria parasites before sequencing. This was done at both genus (*Plasmodium*) and species (P. *falciparum*) levels using photo-induced electron transfer polymerase chain reaction (PET-PCR), which was performed as previously described [19]. Only samples with positive results for both *Plasmodium* genus and *P. falciparum* proceeded to the subsequent step of sequencing.

### Sequencing to detect mutations in drug-resistance genes

Nested PCR was performed to amplify two genes, *pfmdr1* (region 1, codon positions: 86 and 184, and region 2, codon positions: 1034, 1042, and 1246) [20], and the propeller domain of *k13* (codon positions: 430–726) in separate reactions, as previously described [20]. The amplicons from each reaction were visualized on 2% agarose gel stained with RedsafeTM (Biotium, CA, USA). Capillary sequencing was performed using forward and reverse primers with the BIG dye terminator chemistry v3.1 (Applied Biosystems, UK), according to the protocol adopted from CDC in Atlanta, USA [20].

Downstream analysis was done using Geneious® analysis software version 2022.2.2 (Biomatters, New Zealand; www.geneious.com) as described by others [20]. Raw sequence reads were cleaned using geneious default setting, and reads with high-quality scores (the percentage of high-quality bases, ≥70%) were retained for further analysis. The *pfmdr1 and k13* sequences of 3D7 were used as references and single nucleotide polymorphisms (SNPs) detected in one or both strands were considered true SNPs.

### Data management

The SNP data was entered into Microsoft Excel 2016 and later exported into R Studio software (version 4.1.3) for validation, cleaning, and analysis. For the *pfmrd*1 gene, the analysis focused on the three SNPs (N86**<u>Y</u>**, Y184F, and D1246**<u>Y</u>**) and their corresponding haplotypes, which have been associated with reduced susceptibility to lumefantrine. The findings were summarized and presented in text, tables, figures, and maps showing the prevalence and spatial as well as temporal changes of different SNPs and/or haplotypes.

## Results

A total of 2,015 samples were collected in the TES, which were conducted from 2016 to 2021 from subjects receiving AL for the treatment of uncomplicated falciparum malaria. The study that was conducted in 2016 has been published [20], while the 2018 and 2019 studies are available online [21, 22] and others done from 2020 to 2021 have not yet been published. All samples were sequenced for detection of drug resistance mutations in *pfmdr1* and *k13* genes (Table 1). Sequencing success was over 87.8% in both genes, but the *pfmdr1* region 1 had a higher success rate, reaching 94.9% (Table 1).

**Table 1:** Number of samples collected through TES from 2016 to 2021 and sequenced for detection of markers of drug resistance.

| Year | # Samples Sequenced | *k13* * | *pfmdr1*-region 1* | *pfmdr1*-region2* |
| --- | --- | --- | --- | --- |
| 2016 | 423 | 376(88.9 %) | 423(100 %) | 423(100 %) |
| 2018 | 426 | 359(84 %) | 401(94 %) | 379(89 %) |
| 2019 | 403 | 353(88 %) | 371(91.7 %) | 372(92 %) |
| 2020 | 412 | 348(84 %) | 378(93 %) | 330(80 %) |
| 2021 | 351 | 333(95 %) | 335 (95 %) | 270(77 %) |
| **Total** | **2,015** | **1,769(87.8 %)** | **1,908(94.7 %)** | **1,774(88.0 %)** |

#Number of samples that attempted sequenced, and \*number and proportions of samples which passed the sequencing quality score criteria and were therefore used in the analysis.

### Polymorphisms in *k13* gene

Overall, 1,778 (88.2%) samples were successfully sequenced for *k13,* and the majority (1,724/1,769, 97.5%) had wild-type parasites. Of the successfully sequenced samples, 45 (2.5%) had *k13* mutations, with 37 (2.1%) samples carrying synonymous mutations at codons P417P, C469C, R471R, V487V, F505F, G538G, R539R, and S624S. Eight samples (0.4%) had non-synonymous mutations with seven SNPs that are not validated by WHO as molecular markers of ART-R (I416V, E433D, R471S, P475S, A578S, and Q613E). Only one sample from Morogoro in 2020 had R622**I** (a WHO-validated marker of ART-R) in a recurrent infection on day 14 (Table 2).

**Table 2:** *k*13 mutations among samples collected during TES in the eight regions of Tanzania from 2016 to 2021.

| **Year** | **Region** | **#Sequenced** | ***k13*\*** | ***k13*\*\***<br>**(wild-type)** | ***k13*\*\***<br>**(mutant)** | ***k13* \*\***<br>**(SYN**<br>**mutation)** | ***k13*\*\***<br>**(NS**<br>**mutation)** |
| --- | --- | --- | --- | --- | --- | --- | --- |
| **2016** | Kigoma | 118 | 101(85.6 %) | 95(94.1 %) | 6(5.9 %) | 3(2-R471 &<br>C469) | 3(2-Q613E<br>&I416V) |
|  | Morogoro | 104 | 96(92.3 %) | 92(95.8 %) | 4(4.2 %) | 2(R471 &<br>R539) | 2(E433D &<br>A578S) |
|  | Pwani | 92 | 79(85.9 %) | 75(94.9 %) | 4(5.1 %) | 4(P417, C469 &<br>2-V487) | 0 |
|  | Tanga | 109 | 100(91.7 %) | 95(95.0 %) | 5(5.0 %) | 4(P417 & 3-C469) | 1(R471S) |
| **2018** | Kigoma | 111 | 96(86.5 %) | 96(100 %) | 0(0.0 %) | 0 | 0 |
|  | Morogoro | 99 | 87(87.9 %) | 84(96.6 %) | 3(3.4 %) | 3(G538 & 2-R539) | 0 |
|  | Pwani | 97 | 79(81.4 %) | 76(96.2 %) | 3(3.8 %) | 3(S624) | 0 |
|  | Tanga | 119 | 97(81.5 %) | 97(100 %) | 0(0.0 %) | 0 | 0 |
| **2019** | Mbeya | 95 | 90(94.7 %) | 86(95.6 %) | 4(4.4 %) | 4(P417, 2- C469<br>& R539) | 0 |
|  | Mtwara | 93 | 88(94.6 %) | 84(95.5 %) | 4(4.5 %) | 3(C469 & 2-F505) | 1(P475S) |
|  | Tabora | 111 | 96(86.5 %) | 94(98.0 %) | 2(2.0 %) | 2(P417) | 0 |
|  | Mwanza | 104 | 79(76.0 %) | 79(100 %) | 0(0.0 %) | 0 | 0 |
| **2020** | Pwani | 97 | 83(85.7 %) | 82(98.8 %) | 1(1.2 %) | 1(R539) | 0 |
|  | Kigoma | 103 | 70(68.0 %) | 66(94.3 %) | 4(5.7 %) | 4(P417) | 0 |
|  | Morogoro | 96 | 95(90.0 %) | 91(95.8 %) | 4(4.2 %) | 3(P417) | 1(R622I) |
|  | Tanga | 116 | 100(86.2 %) | 100(100 %) | 0(0.0 %) | 0 | 0 |
| **2021** | Mbeya | 48 | 43(89.6 %) | 43(100 %) | 0(0.0 %) | 0 | 0 |
|  | Mtwara | 87 | 87(100 %) | 87(100 %) | 0(0.0 %) | 0 | 0 |
|  | Mwanza | 106 | 96(90.6 %) | 96(100 %) | 0(0.0 %) | 0 | 0 |
|  | Tabora | 110 | 107(97.3 %) | 106(99.1 %) | 1(0.9 %) | 1(Y558) | 0 |
| **Total** |  | **2,015** | **1,769(87.8 %)** | **1,724(97.5 %)** | **45(2.5 %)** | **37(2.1 %)** | **8(0.4 %)** |

*k13* = *P. falciparum* kelch 13, SYN = synonymous mutations, and NS = non-synonymous mutations
\*number of samples that passed the sequencing quality score criteria; \*\*number of samples with either wild-type or mutant genotypes and their
corresponding proportions where appropriate.

### Polymorphisms and haplotypes in the *pfmdr1* gene

The number of successfully sequenced samples for the *pfmdr1* regions covering the three mutations were 1,912 (94.9%) for codons 86 and 184, and 1,774 (88.0%) for codon 1246 (Table 1 and Table 3). For *pfmdr1* codon 86, all except two samples (which had 86**Y** mutations in 2016) carried N86 (wildtype). Mutations at Y184**F** were more prevalent and increased from 33.9% in 2016 to about 60.5% in 2021 (Figure 2). D1246**Y** mutations occurred in four samples (0.2%): two from Morogoro and Tanga in 2016 and two other mutations from Kigoma and Morogoro in 2020. The *pfmdr1* haplotypes (N86**Y,** Y184**F,** and D1246**Y**) were constructed with 1,711 (85.0%) samples, and 985 (57.6%) of these had NYD, 720 (42.1%) had N**F**D, while six samples (0.4%) had minor haplotypes (three with NY**Y** = 0.2%, **YF**D in two, 0.1%, and N**F**Y in one sample 0.1%). Between 2016 and 2021, *pfmdr1* haplotype NYD showed a decrease of from 66.1% to 45.2% (Figure 3) while N**F**D increased from 38.5% to 54.7% (Figure 4), but these changes varied among the study regions.

**Figure 2:**
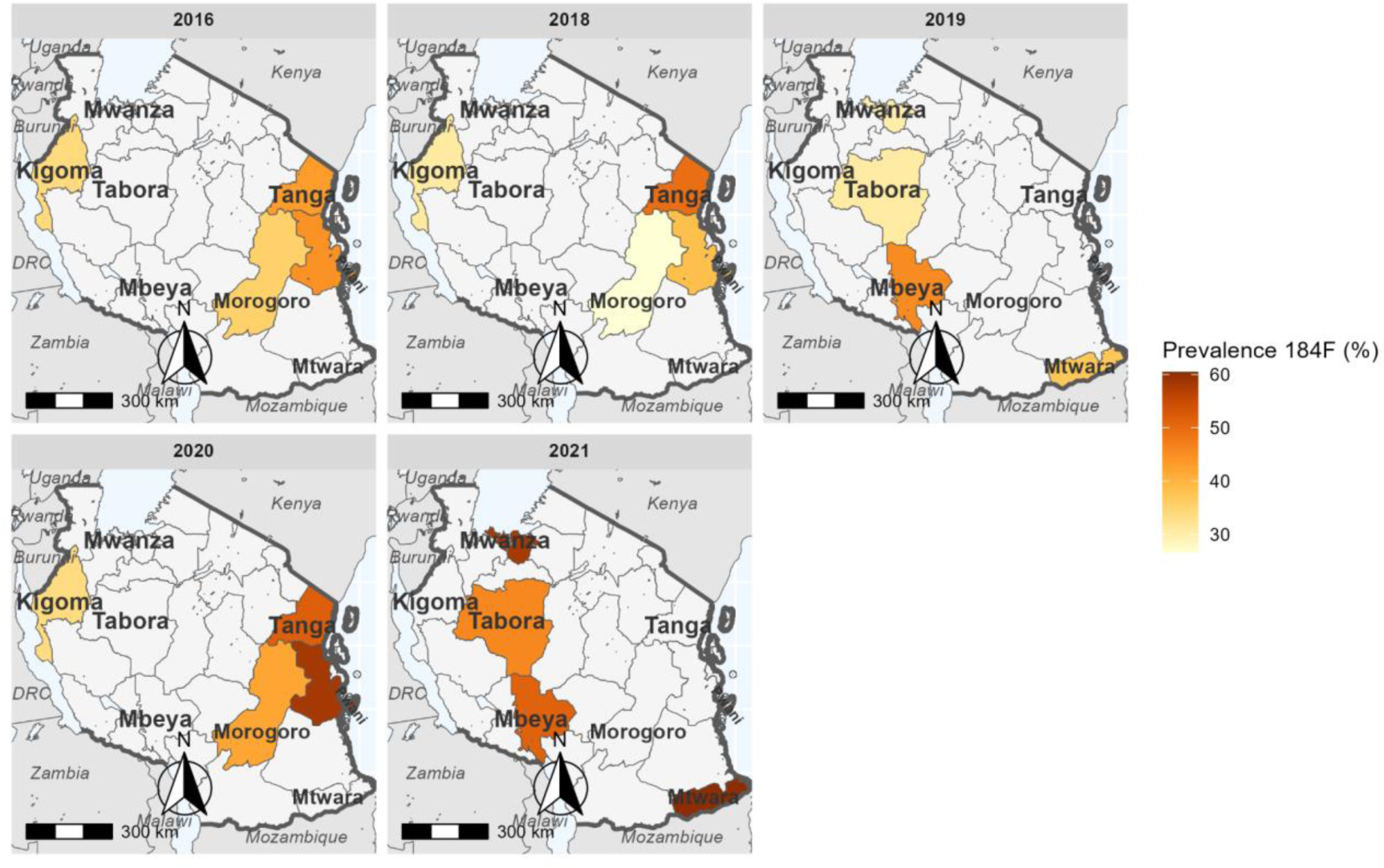
Trend of 184F mutation in the eight regions of Mainland Tanzania from 2016 to 2021.

**Figure 3:**
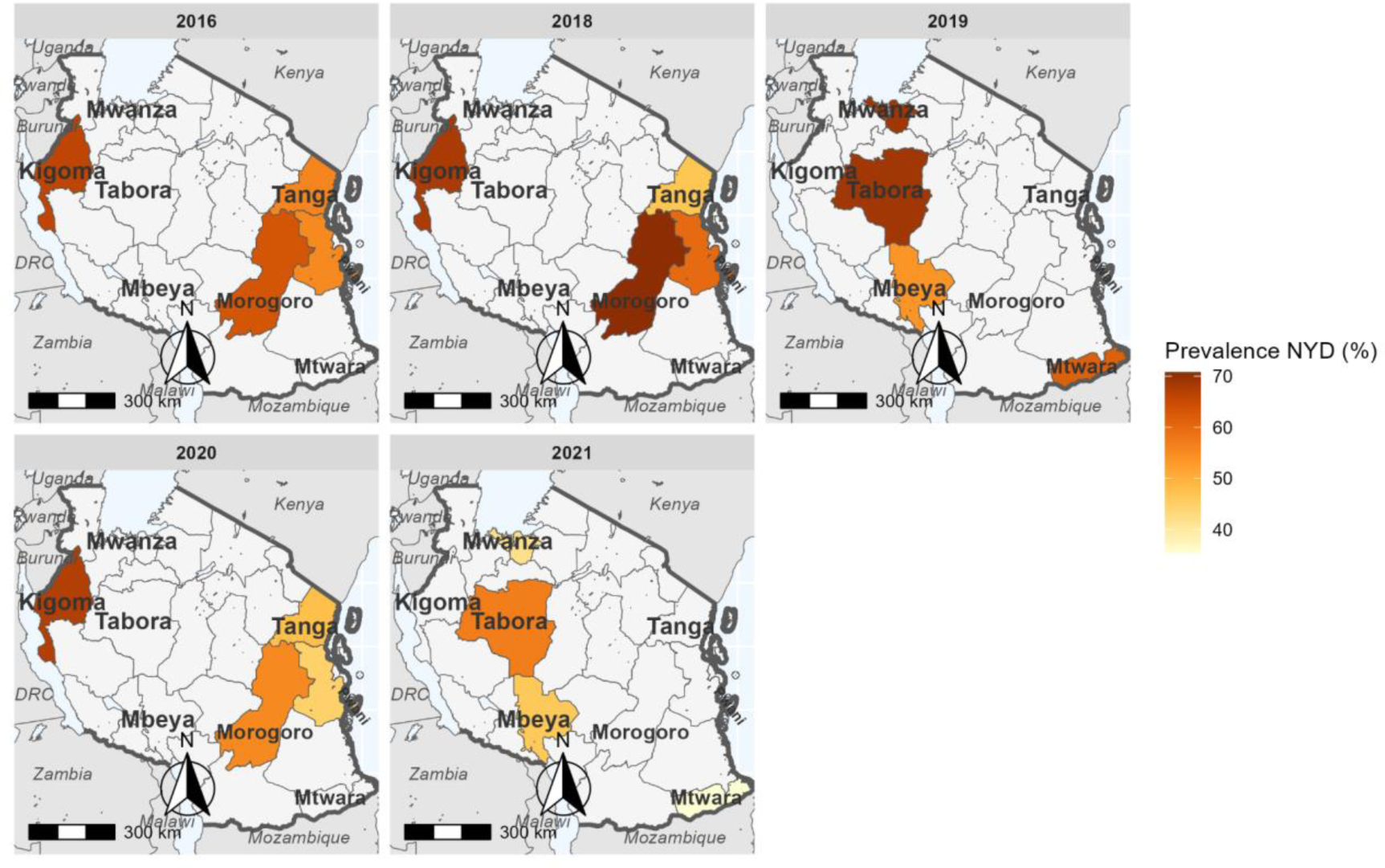
Trend of *pfmdr*1 NYD haplotype in the eight regions of Mainland Tanzania from 2016 to 2021.

**Figure 4:**
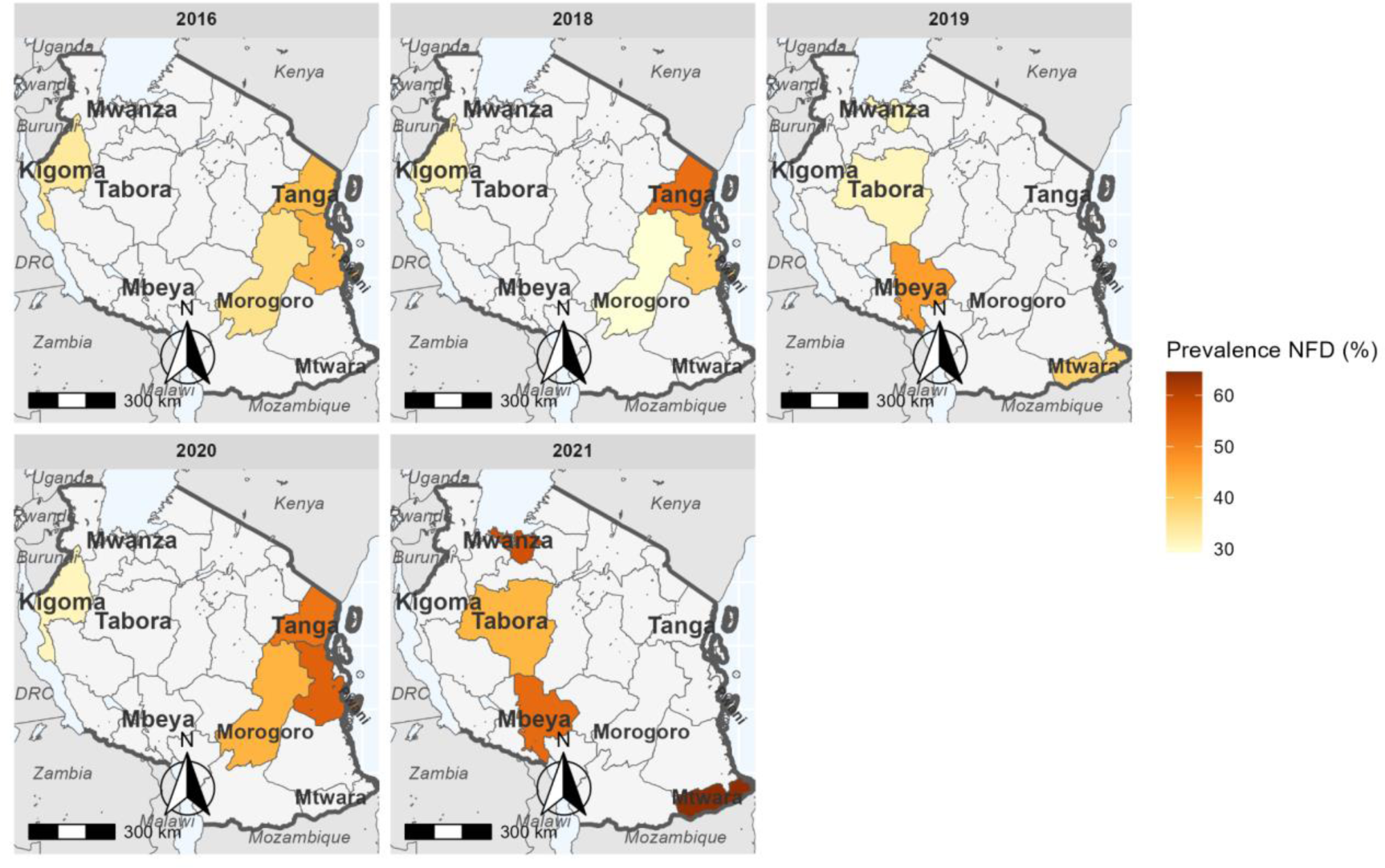
Trend of *pfmdr*1 N**F**D haplotype in the eight regions of Mainland Tanzania from 2016 to 2021.

**Table 3:**
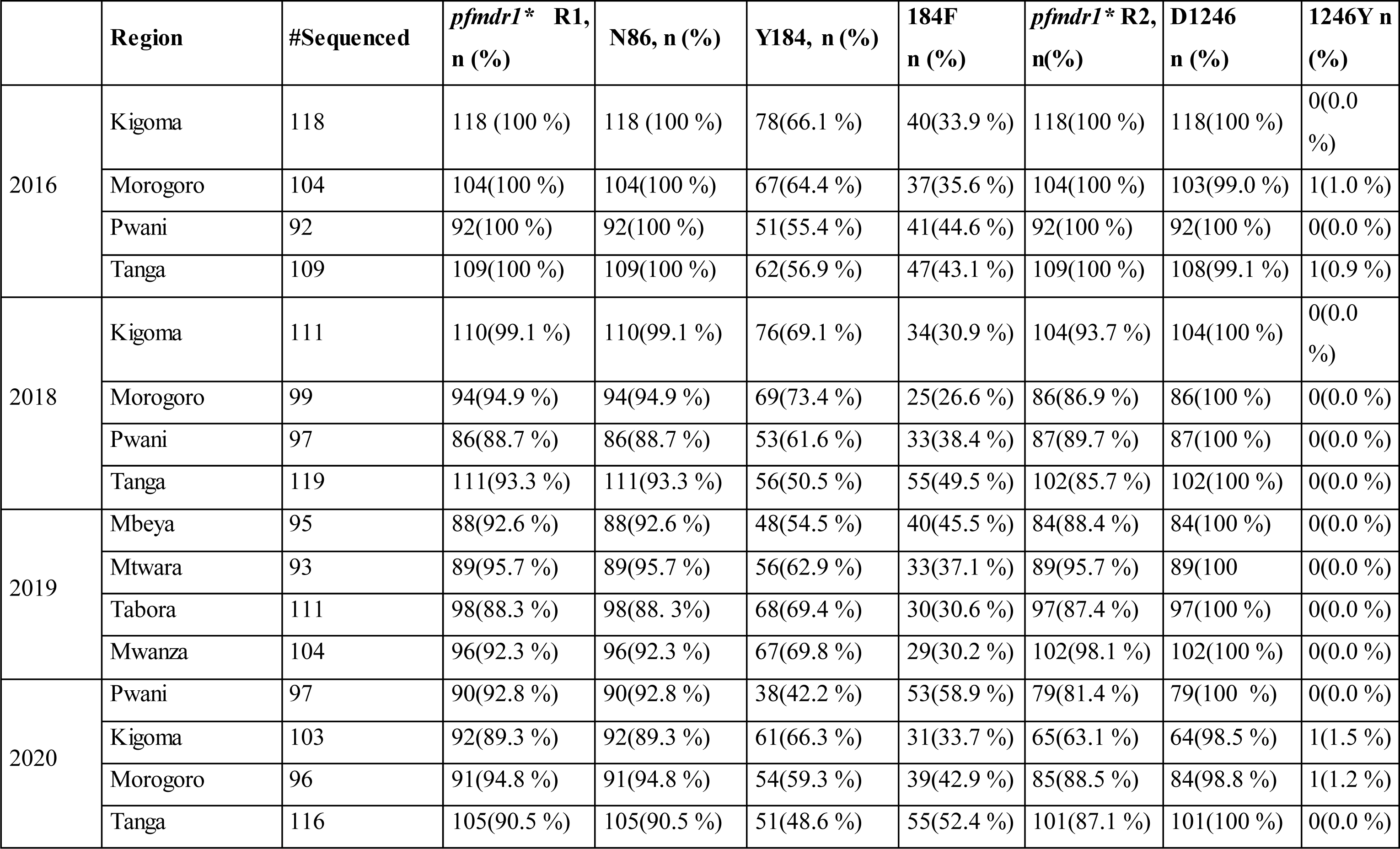
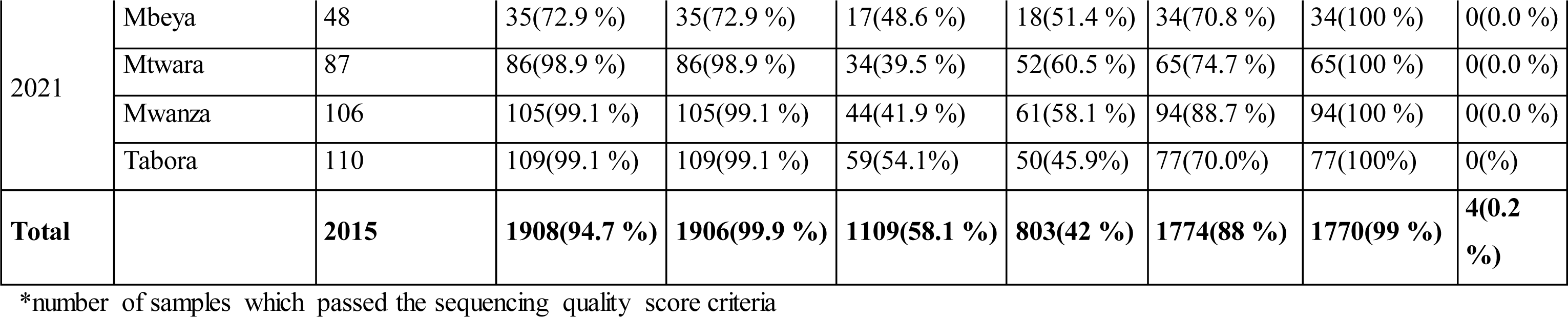
*pfmdr1* mutations among samples collected in the eight regions from 2016 to 2021.

|  | **Region** | **#Sequenced** | ***pfmdr1*\* R1,**<br>**n (%)** | **N86, n (%)** | **Y184, n (%)** | **184F**<br>**n (%)** | ***pfmdr1*\* R2,**<br>**n(%)** | **D1246**<br>**n (%)** | **1246Y n**<br>**(%)** |
| --- | --- | --- | --- | --- | --- | --- | --- | --- | --- |
| 2016 | Kigoma | 118 | 118 (100 %) | 118 (100 %) | 78(66.1 %) | 40(33.9 %) | 118(100 %) | 118(100 %) | 0(0.0 %) |
|  | Morogoro | 104 | 104(100 %) | 104(100 %) | 67(64.4 %) | 37(35.6 %) | 104(100 %) | 103(99.0 %) | 1(1.0 %) |
|  | Pwani | 92 | 92(100 %) | 92(100 %) | 51(55.4 %) | 41(44.6 %) | 92(100 %) | 92(100 %) | 0(0.0 %) |
|  | Tanga | 109 | 109(100 %) | 109(100 %) | 62(56.9 %) | 47(43.1 %) | 109(100 %) | 108(99.1 %) | 1(0.9 %) |
| 2018 | Kigoma | 111 | 110(99.1 %) | 110(99.1 %) | 76(69.1 %) | 34(30.9 %) | 104(93.7 %) | 104(100 %) | 0(0.0 %) |
|  | Morogoro | 99 | 94(94.9 %) | 94(94.9 %) | 69(73.4 %) | 25(26.6 %) | 86(86.9 %) | 86(100 %) | 0(0.0 %) |
|  | Pwani | 97 | 86(88.7 %) | 86(88.7 %) | 53(61.6 %) | 33(38.4 %) | 87(89.7 %) | 87(100 %) | 0(0.0 %) |
|  | Tanga | 119 | 111(93.3 %) | 111(93.3 %) | 56(50.5 %) | 55(49.5 %) | 102(85.7 %) | 102(100 %) | 0(0.0 %) |
| 2019 | Mbeya | 95 | 88(92.6 %) | 88(92.6 %) | 48(54.5 %) | 40(45.5 %) | 84(88.4 %) | 84(100 %) | 0(0.0 %) |
|  | Mtwara | 93 | 89(95.7 %) | 89(95.7 %) | 56(62.9 %) | 33(37.1 %) | 89(95.7 %) | 89(100 %) | 0(0.0 %) |
|  | Tabora | 111 | 98(88.3 %) | 98(88.3 %) | 68(69.4 %) | 30(30.6 %) | 97(87.4 %) | 97(100 %) | 0(0.0 %) |
|  | Mwanza | 104 | 96(92.3 %) | 96(92.3 %) | 67(69.8 %) | 29(30.2 %) | 102(98.1 %) | 102(100 %) | 0(0.0 %) |
| 2020 | Pwani | 97 | 90(92.8 %) | 90(92.8 %) | 38(42.2 %) | 53(58.9 %) | 79(81.4 %) | 79(100 %) | 0(0.0 %) |
|  | Kigoma | 103 | 92(89.3 %) | 92(89.3 %) | 61(66.3 %) | 31(33.7 %) | 65(63.1 %) | 64(98.5 %) | 1(1.5 %) |
|  | Morogoro | 96 | 91(94.8 %) | 91(94.8 %) | 54(59.3 %) | 39(42.9 %) | 85(88.5 %) | 84(98.8 %) | 1(1.2 %) |
|  | Tanga | 116 | 105(90.5 %) | 105(90.5 %) | 51(48.6 %) | 55(52.4 %) | 101(87.1 %) | 101(100 %) | 0(0.0 %) |
| 2021 | Mbeya | 48 | 35(72.9 %) | 35(72.9 %) | 17(48.6 %) | 18(51.4 %) | 34(70.8 %) | 34(100 %) | 0(0.0 %) |
|  | Mtwara | 87 | 86(98.9 %) | 86(98.9 %) | 34(39.5 %) | 52(60.5 %) | 65(74.7 %) | 65(100 %) | 0(0.0 %) |
|  | Mwanza | 106 | 105(99.1 %) | 105(99.1 %) | 44(41.9 %) | 61(58.1 %) | 94(88.7 %) | 94(100 %) | 0(0.0 %) |
|  | Tabora | 110 | 109(99.1 %) | 109(99.1 %) | 59(54.1%) | 50(45.9%) | 77(70.0%) | 77(100%) | 0(%) |
| **Total** |  | **2015** | **1908(94.7 %)** | **1906(99.9 %)** | **1109(58.1 %)** | **803(42 %)** | **1774(88 %)** | **1770(99 %)** | **4(0.2 %)** |

\*number of samples which passed the sequencing quality score criteria

## Discussion

Antimalarial resistance threatens the effectiveness of current malaria treatments much like it has done for various antimalarials over the past half a century [1]. In recent years, reports of ART-R in Africa [1] have been a growing concern because of the potential emergence and spread of ACTs resistance. Thus, there is a critical need to monitor the effectiveness of these strategies using different approaches, such as TES and MMS. Tracking the emergence and spread of molecular markers of resistance to artemisinin and partner drugs is crucial so as to maintain the effectiveness of malaria treatment, prevent the spread of resistant strains and contribute to global efforts to respond to ART-R as well as control and eliminate malaria. The current study aimed to assess the trends of molecular markers of drug resistance in two genes, *k13* and *pfmdr1,* using the data collected in Mainland Tanzania between 2016 and 2021. Over the study period, WHO-validated mutations in *k13*, which are associated with ART-R, were only detected in one sample, while significant changes occurred in the *pfmdr1* gene.

This study reported less than 2.5% of non-synonymous mutations in *k13* gene, and only one sample had validated *k*13 mutation (R622**I**) from Morogoro in 2020. The findings align with most reports in some African countries in which no or a very low prevalence of *k13* mutations has been reported [23,24]. Until recently, R622**I** has been reported in three countries: Ethiopia, Eritrea, and Sudan [25], and this is the first report of the R622**I** mutation in Tanzania. In Eritrea, an increase in R622**I** prevalence from less than 10% in 2016 to 20% in 2019 has been reported [7], while in Ethiopia, the mutation has been reported in several studies, with prevalence ranging from 2.4% to 9.8% [8, 9]. Similarly, it has been shown that parasites with R622**I** mutations tend also to carry deletions of histidine-rich protein 2/3 (*hrp2/3)* gene, which is linked to the failure of malaria rapid diagnostic tests that detect the HRP2 antigen to detect *P. falciparum* infections. In a recent Tanzania-wide surveillance, the R622**I** was also detected in one sample from Njombe region in 2021, in an area close to Morogoro where the 2020 sample was detected [10]. Njombe is a region that reported more parasites with *hrp2/3* single gene deletions in 2021, compared to other regions of Tanzania [26]. More studies will be needed to explore if the parasites with R622**I** are co-emerging with the gene deletions as reported in Ethiopia and Eritrea.

Despite the reports of ART-R in Rwanda [9] and Uganda [27], this study did not report any other validated mutations in these study sites in Tanzania apart from R622**I**. However, previous studies in Tanzania reported the presence of R561**H** mutation in two regions of Pwani in 2020 [28] and Geita [29], and this analysis revealed no evidence for presence of these mutations in these study sites during this period. Recently, the R56I**H** mutation was detected at high prevalence in some parts of Kagera (reaching over 20%), while few samples with the same mutations were also detected in Tabora, Njombe and Manyara in 2021 [10]. More surveys in Kagera have observed an increase in these mutations in some districts and a spread from three districts in 2021 to five in 2023 (Ishengoma et al., unpublished data). Another mutation (A675V) has also been reported in Kagera, but at low prevalence compared to R561**H,** suggesting a potential threat of spreading ART-R in Kagera and other regions (Ishengoma, unpublished data). Therefore, there is an urgent need for other strategies apart from TES for effectively monitoring the spread of ART-R in Tanzania because the presence of these mutations could not be captured by TES in the studies that were done before and after deployment of ACTs in 2006.

The polymorphisms in the *pfmdr*1 gene have been associated with several anti-malarial drugs, including lumefantrine, CQ, and AQ. The N86 and D1246 alleles were observed to be near fixation, and N86, which has been linked to decreased susceptibility to lumefantrine and increased susceptibility to AQ, was only detected in two samples in 2016 (in Pwani and Tanga). Studies from various parts of Africa have also reported similar results, and this has been linked to AL selecting the N86 allele [30–32]. There was a significant increase in both the 184**F** and N**F**D haplotypes over the years, and this might be linked to selection caused by lumefantrine. Another study conducted in Bagamoyo districts in Tanzania using the samples collected from 2006 to 2011 reported similar findings, with a high prevalence of 184**F** mutations, and an increase from 14.0% in 2006 to 35.0% in 2011 [13]. Several other studies conducted in Africa have also documented an increase of 184**F**, leading to a rise in N**F**D haplotype, possibly due to the continued use of AL as a first-line treatment. In Tanzania, the trends of both 184**F** and N**F**D haplotype across regions appear to be homogeneous, indicating the drug selection pressure might be similar throughout the country. Similar findings have been reported in other sub-Saharan African countries, where temporal trends of the Y184**F** mutations and N**F**D haplotypes have been associated with reduced susceptibility to the lumefantrine component of AL [33,34]. It is important to note that the changes in *pfmdr1* markers were not associated with reduced efficacy of AL, which was >95.0% for all years [18]. Although these mutations have not been linked to ACT failure, it’s essential to monitor their spread and possible association (together with other markers) with ACT resistance in different endemic countries to inform malaria case management strategies.

## Conclusions

This is the first report of the R622I (k13 validated mutation) in Tanzania. N86 wildtype, which is associated with decreased susceptibility to lumefantrine and increased susceptibility to AQ and CQ, is near-fixation together with D1246. Changes were observed in *pfmdr1*, with an increase in Y184**F** mutations and N**F**D haplotype reaching over 50% in all regions except Tabora, with over 42% in 2021. Despite the reports of ART-R in Rwanda and Uganda, this study did not report any other validated mutations in these study sites in Tanzania apart from R622I. Following detection of ART-R (561**H**) in the Kagera region, which was not captured by TES, intensified molecular surveillance is urgently needed to monitor the trends of drug resistance markers and their potential impact on the performance of ACTs.

## List of abbreviations

ACT: artemisinin-based combination therapy
AL: artemether-lumefantrine
ASAQ: artesunate + amodiaquine
AQ: Amodiaquine
CDC: US Centers for Disease Control and Prevention
CQ: chloroquine
DBS: dried blood spot
DNA: deoxyribonucleic acid
DP: Dihydroartemisinin–Piperaquine
MRCC: Medical Research Coordinating Committee of NIMR
NIMR: National Institute for Medical Research
NMCP: National Malaria Control Programme
PARMA: Partnership for Antimalarial Resistance Monitoring in Africa network
k13: *Plasmodium falciparum* Kelch 13 gene
Pfmdr1: *Plasmodium falciparum* multi-drug resistance 1 gene
PMI: US President’s Malaria Initiative
PCR: polymerase chain reaction
SNP: single nucleotide polymorphism
SP: sulphadoxine-pyrimethamine
SSA: sub-Saharan African
TES: therapeutic efficacy study
TWG: Technical Working Group
USA: United States of America
WHO: World Health Organization

## Ethical approval

All the studies which generated the samples obtained ethical clearance from the Medical Research Coordinating Committee (MRCC) of the National Institute for Medical Research (NIMR - MRCC) in Tanzania.

## Permission to publish

Permission to publish the manuscript was sought and provided by the Director General of NIMR.

## Availability of data and materials

The datasets generated and/or analyzed during the current study are available from the corresponding author upon reasonable request.

## Authors’ contributions

DSI, CB, RAM, MDS and CIM conceived of the study, designed the experiments and took part in the laboratory analysis of samples with the support of ET, NWL, MV, LM, ESH and UV. DSI, CIM, BN, EK, MA, FF, SB, MC, MKM, RAK and FM implemented TES in the 8 regions with the support of SM, RM, FM, FC, DB, RN, MW, SN, KB, NS, CK and ER. RAM, CB and MDS generated and analyzed the data under the supervision of DSI: CB and DSI wrote the manuscript. All authors reviewed and approved the final manuscript.

## Acknowledgments

The authors are indebted to the children and their parents/guardians for agreeing to take part in the studies and attending the follow-up visits despite the long durations of follow-up. They extend their appreciation to health facilities’ staff, stakeholders, and colleagues from implementing partners and local health authorities for their support during the entire period of implementing the TES and this study. The support provided by the regional and district authorities is greatly acknowledged. Technical and logistic support provided by the CDC, USAID/PARMA, RTI, and NIMR teams is highly appreciated. Permission to publish this paper has been granted by the Director General of NIMR.

## Disclaimer

The findings and conclusions presented in this report are those of the authors and do not necessarily reflect the official position of the Centers for Disease Control and Prevention and of the US Agency for International Development.

